# Non-cyanobacterial diazotrophs mediate dinitrogen fixation in biological soil crusts during early crust formation

**DOI:** 10.1101/013813

**Authors:** Charles Pepe-Ranney, Chantal Koechli, Ruth Potrafka, Cheryl Andam, Erin Eggleston, Ferran Garcia-Pichel, Daniel H Buckley

## Abstract

Biological soil crusts (BSC) are key components of ecosystem productivity in arid lands and they cover a substantial fraction of the terrestrial surface. In particular, BSC N_2_-fixation contributes significantly to the nitrogen (N) budget of arid land ecosystems. In mature crusts, N_2_-fixation is largely attributed to heterocystous cyanobacteria, however, early successional crusts possess few N_2_-fixing cyanobacteria and this suggests that microorganisms other than cyanobacteria mediate N_2_-fixation during the critical early stages of BSC development. DNA stable isotope probing (DNA-SIP) with ^15^N_2_ revealed that *Clostridiaceae* and *Proteobacteria* are the most common microorganisms that assimilate ^15^N_2_ in early successional crusts. The *Clostridiaceae* identified are divergent from previously characterized isolates, though N_2_fixation has previously been observed in this family. The Proteobacteria identified share >98.5 %SSU rRNA gene sequence identity with isolates from genera known to possess diazotrophs (e.g. *Pseudomonas*, *Klebsiella*, *Shigella*, and *Ideonella*). The low abundance of these heterotrophic diazotrophs in BSC may explain why they have not been characterized previously. Diazotrophs play a critical role in BSC formation and characterization of these organisms represents a crucial step towards understanding how anthropogenic change will affect the formation and ecological function of BSC in arid ecosystems.

## 2 INTRODUCTION

Biological soil crusts (BSC) are specialized microbial communities that form at the soil surface in arid environments and they fill a variety of important ecological functions. BSCs occupy plant interspaces and cover a wide, global geographic range (Garcia-Pichel *et al.*, 2003). For example, in some regions on the Colorado Plateau BSCs cover 80% of the ground (Karnieli *et al.*, 2003). The global biomass of BSC cyanobacteria alone is estimated at 54 × 10^12^ g C (Garcia-Pichel *et al.*, 2003). BSD nitrogen fixation (N_2_-fixation) is responsible for significant input of nitrogen (N) to arid environments (Evans and Belnap, 1999; Belnap, 2003). Interestingly, much of this fixed N is exported from the crusts in dissolved form through percolation or runoff and little is lost to volatilization (Johnson *et al.*, 2007). The presence of BSC is positively correlated with vascular plant survival due in part to N inputs from BSC (for review of BSC-vascular plant interactions see Belnap *et al.* 2003). These microbial ecosystems are not immune to climate change and changes in precipitation and temperature could alter BSC microbial community structure/membership and possibly BSC diazotroph diversity and N_2_-fixation (Garcia-Pichel *et al.*, 2013).

BSC are highly susceptible to natural and anthropogenic disturbance (Garcia-Pichel *et al.*, 2013) Succession in BSC communities is characterized by transition from early successional “light” crusts to mature “dark” crusts (Belnap, 2002; Yeager *et al.*, 2004). Motile nonheterocystous cyanobacteria(e.g.*Microcoleus vaginatus* or *M. steenstrupii*), which cannot fix N_2_ are pioneer colonizers of early successional crusts and are abundant in all types of BSCs (Garcia-Pichel *et al.*, 2013). Successional development of mature crust is accompanied by a change in color produced by secondary colonization with non-motil N_2_-fixing heterocystous cyanobacteria which produce sunscreen compounds that reduce soil albedo (Belnap, 2002; Yeager *et al.*, 2004). These heterocystous cyanobacteria (e.g. *Scytonema*, *Spirirestis*, and *Nostoc*) increase in abundance during crust development and are more abundant in mature crusts (Yeager *et al.*, 2007, 2012) Heterocystous cyanobacteria are numerically dominant in surveys of BSC *nifH* gene diversity (Yeager *et al.*, 2004, 2007, 2012). For example, 89 percent of 693 *nifH* sequences derived from Colorado Plateau and New Mexico BSC were attributed to heterocystous cyanobacteria (Yeager *et al.*, 2007). Other BSC *nifH* sequences are attributed to *Alpha-*, *Beta-*, and *Gammaproteobacteria*, as well as a *nifH* clade (*nifH* cluster III) that includes diverse anaerobes such as clostridia, sulfate reducing bacteria, and anoxygenic phototrophs (Steppe *et al.*, 1996; Yeager *et al.*, 2007).

Two lines of evidence suggest that nitrogen fixers other than phototrophs are important in early-successional crusts. First, the contributions of early successional BSC to N_2_-fixation in arid ecosystems may have been systematically under-estimated. The high abundance of heterocystous cyanobacteria at the surface of mature crusts, where acetylene reduction assay rates are often maximal, is generally taken as evidence that BSC N_2_-fixation occurs primarily in mature crusts and is dominated by heterocystous cyanobacteria. However, rates of BSC N_2_-fixation are typically determined by areal measurements made at the crust surface with the acetylene reduction assay and vary significantly across samples and studies (Evans and Lange, 2001). The reasons for inter-site and inter-study variability are complex and likely include the spatial heterogeneity of BSC (Evans and Lange, 2001). The acetylene reduction assay is also subject to methodological artifacts that can complicate comparisons between samples that differ in their physical and biological characteristics (see Belnap 2001 for review). In particular, N_2_-fixation in early successional BSC is maximal below the crust surface (Johnson *et al.*, 2005) and hence diffusional limitation (of both acetylene and ethylene) across the crust surface can cause severe underestimates if they do not allow for sufficiently long incubation times (Johnson *et al.*, 2005). If BSC N_2_-fixation is instead estimated by integrating rates across a depth profile (which eliminates constraints from diffusional limitation), then total rates of N_2_-fixation do not differ significantly between early successional and mature BSC (Johnson *et al.*, 2005). This result suggests that diazotrophs other than heterocystous cyanobacteria may be important contributors to N_2_-fixation in early successional BSC communities as early successional BSC possess few heterocystous cyanobacteria and these are present near the crust surface. Second, the bare soils that are colonized during the process of early crust formation are unconsolidated and oligotrophic in many respects, with much lower N content than adjacent crusts (Beraldi-Campesi *et al.*, 2009), and the cyanobacteria that are typical colonization pioneers (*Microcoleus* spp., (Johnson *et al.*, 2005) Garcia-Pichel and Wojciechowski 2009), are unable to fix nitrogen as they lack that genetic capacity (Starkenburg *et al.*, 2011; Rajeev *et al.*, 2013).

To determine the agency of nitrogen fixation in early developmental crusts, we conducted ^15^N_2_ DNA stable isotope probing (DNA-SIP) experiments with early successional Colorado Plateau BSC conspicuously devoid of significant surface populations of heterocystous cyanobacteria. DNA-SIP with ^15^N_2_ has not been previously attempted with BSC. DNA-SIP provides an accounting of *active* diazotrophs on the basis of ^15^N_2_ assimilation into DNA whereas *nifH* clone libraries merely account for microbes with the genomic potential for N_2_-fixation. Further, we investigate the distribution of these active diazotrophs in surveys of microbial diversity conducted on BSC over a range of spatial scales and soil types (Garcia-Pichel *et al.*, 2013; Steven *et al.*, 2013).

## 3 MATERIALS AND METHODS

### 3.1 BSC SAMPLING AND INCUBATION CONDITIONS

BSC samples were taken from the Green Butte site near Moab, Utah as previously described (site “CP3”; latitude N 38°44’55.1”, longitude W 109°44’37.1”; Beraldi-Campesi *et al.* 2009). All samples were from early successional ‘light’ crusts as described by (Johnson *et al.*, 2005). Early successional BSC samples (37.5 cm^2^, average mass 35 g) were incubated in sealed chambers under controlled atmosphere and in 16 h light / 8 h dark for 4 days. Crusts were sampled and transported while dry and wetted at initiation of the experiment. Water was added to each sample to fully saturate the soil, but avoid visible ponding. The samples were then placed in air-tight sealed incubation containers for the rest of the experiment, so that soil and atmosphere remained saturated through the incubation period. The water was amended with calcium bicarbonate to yield a final concentration of 3 mM, so that autotrophy could proceed unimpeded. The control treatment received a headspace of air and the experimental treatment received a headspace containing ^15^N_2_ (>98% atom ^15^N_2_). ^15^N_2_ (100%) gas was purchased from Sigma-Aldrich (St. Louis, MO). We used a composition of 75% ^15^N_2_ in helium for the initial incubation headspace. Four crust samples were treated and incubated (two control and two experimental). One control/experimental crust pair was collected at day 2 and the other at day 4. Acetylene reduction rates were measured daily. Acetylene reduction rates increased over the course of the experiment (0.8, 4.8, 8.8, and 14.5 *μ*moles m-2 hr−1 ethylene for days 1 through 4, respectively).

### 3.2 DNA EXTRACTION

DNA was extracted for DNA-SIP at 2 and 4 days. DNA was extracted from 1 g of BSC. DNA from each sample was extracted using a MoBio (Carlsbad, CA) UltraClean Mega Soil DNA Isolation Kit (following manufacturer’s protocol, but lysis was done as previously described (Strauss *et al.*, 2011)), and then gel purified to select high molecular weight DNA (>4 kb) using a 1% low melt agarose gel and *β*-agarase I for digestion (manufacturer’s protocol, New England Biolabs, M0392S). Extracts were quantified using PicoGreen nucleic acid quantification dyes (Molecular Probes).

### 3.3 FORMATION OF CSCL EQUILIBRIUM DENSITY GRADIENTS

CsCl gradient fractionation was used to separate the DNA into 36 gradient fractions on the basis of buoyant density. CsCl density gradients were formed in 4.7 mL polyallomer centrifuge tubes filled with gradient buffer (15 mM Tris-HCl, pH 8; 15 mM EDTA; 15 mM KCl) which contained 1.725 g mL-1 CsCl. CsCl density was checked with a digital refractometer as described below. A total of 2.5-5.0 *μ*g of DNA was added to each tube, and the tubes mixed, prior to centrifugation. Centrifugation was performed in a TLA-110 fixed angle rotor (Beckman Coulter) at 20°C for 67 hours at 55,000 rpm. (Buckley *et al.*, 2007). Centrifuged gradients were fractionated from bottom to top in 36 equal fractions of 100 *μ*L, using a syringe pump as described previously (Buckley *et al.*, 2007). The density of each fraction was determined using an AR200 refractometer modified to accommodate 5 *μ*L samples as described previously (Buckley *et al.*, 2007). DNA in each fraction was desalted on a filter plate (PALL, AcroPrep Advance 96 Filter Plate, Product Number 8035), using four washes with 300 *μ*L TE per fraction. After each wash, the filter plate was centrifuged at 500 × g for 10 minutes, with a final spin of 20 minutes. Purified DNA from each fraction was resuspended in 50 *μ*L of TE buffer.

### 3.4 PCR, LIBRARY NORMALIZATION AND DNA SEQUENCING

To characterize the distribution of SSU rRNA genes across density gradients, SSU rRNA gene amplicons were generated from 20 gradient fractions per gradient for both unlabeled controls and ^15^N_2_ labeled samples. The 20 fractions analyzed are those expected to contain DNA (both labeled and unlabeled) having buoyant density in the range of 1.66 g mL-1 to 1.77 g mL-1. Barcoded PCR of bacterial and archaeal SSU rRNA genes was carried out using primer set 515F/806R (Walters *et al.*, 2011) (primers purchased from Integrated DNA Technologies). The primer 806R contained an 8 bp barcode sequence, a “TC” linker, and a Roche 454 B sequencing adapter, while the primer 515F contained the Roche 454 A sequencing adapter. Each 25 *μ*L reaction contained 1x PCR Gold Buffer (Roche), 2.5 mM MgCl_2_, 200 *μ*M of each of the four dNTPs (Promega), 0.5 mg mL-1 BSA (New England Biolabs), 0.3 *μ*M of each primers, 1.25 U of Amplitaq Gold (Roche), and 8 *μ*L of template. Each sample was amplified in triplicate. Thermal cycling occurred with an initial denaturation step of 5 minutes at 95°C, followed by 40 cycles of amplification (20 s at 95°, 20 s at 53°, 30 s at 72°), and a final extension step of 5 min at 72C. Triplicate amplicons were pooled and purified using Agencourt AMPure PCR purification beads, following manufacturer’s protocol. Once purified, amplicons were quantified using PicoGreen nucleic acid quantification dyes (Molecular Probes) and pooled together in equimolar amounts. Samples were sent to the Environmental Genomics Core Facility at the University of South Carolina (now Selah Genomics) where they were run on a Roche FLX 454 pyrosequencing machine (FLX-Titanium platform).

### 3.5 DATA ANALYSIS

#### 3.5.1 Sequence quality control

Sequences were initially screened by maximum expected errors at a specific read length threshold (Edgar, 2013) and this has been shown to be as effective as denoising with respect to removing pyrosequencing errors. Specifically, reads were first truncated to 230 nucleotides (nt) (all reads shorter than 230 nt were discarded) and any read that exceeded a maximum expected error threshold of 1.0 was removed. After truncation and max expected error trimming, 91% of original reads remained. Forward primer and barcode were then removed from the high quality, truncated reads. Remaining reads were taxonomically annotated using the “UClust” taxonomic annotation framework in the QIIME software package (Caporaso *et al.*, 2010; Edgar, 2010) with cluster seeds from Silva SSU rRNA database (Pruesse *et al.*, 2007) 97% sequence identity OTUs as reference (release SSU Ref 111). Reads annotated as “Chloroplast”, “Eukaryota”, “Archaea”, “Unassigned” or “mitochondria” were removed from the dataset. Finally, reads were aligned to the Silva reference alignment provided by the Mothur software package (Schloss *et al.*, 2009) using the Mothur NAST aligner (DeSantis *et al.*, 2006). All reads that did not align to the expected amplicon region of the SSU rRNA gene were discarded. Quality control parameters removed 34,716 of 258,763 raw reads. Raw sequences have been uploaded to MG-RAST (MG-RAST ID 4603397.3).

#### 3.5.2 Sequence clustering

Sequences were distributed into OTUs using the UPARSE methodology (Edgar, 2013). Specifically, OTU centroids (i.e. seeds) were identified using USEARCH on non-redundant reads sorted by count. The sequence identity threshold for establishing a new OTU centroid was 97%. After initial OTU centroid selection, select SSU rRNA gene sequences from Yeager *et al.* (2007) were added to the centroid collection. Specifically, Yeager *et al.* (2007) Colorado Plateau or Moab, Utah sequences were added which included the SSU rRNA gene sequences for *Calothrix* MCC-3A (accession DQ531700.1), *Nostoc commune* MCT-1 (accession DQ531903), *Nostoc commune* MFG-1 (accession DQ531699.1), *Scytonema hyalinum* DC-A (accession DQ531701.1), *Scytonema hyalinum* FGP-7A (accession DQ531697.1), *Spirirestis rafaelensis* LQ-10 (accession DQ531696.1). Original centroid sequences that matched selected Yeager *et al.* (2007) (above) sequences with greater than to 97% sequence identity were subsequently removed from the centroid collection. With USEARCH/UPARSE, potential chimeras are identified during OTU centroid selection and are not allowed to become cluster centroids effectively removing chimeras from the read pool. All quality controlled reads were then mapped to cluster centroids at an identity threshold of 97% again using USEARCH. A total of 95.6% of quality controlled reads could be mapped to centroids. Unmapped reads do not count towards sample counts and were removed from downstream analyses. The USEARCH software version for cluster generation was 7.0.1090. Garcia-Pichel *et al.* (2013) and Steven *et al.* (2013) sequences were quality screened by alignment coordinates (described above) and included as input to USEARCH for OTU centroid selection and subsequent mapping to OTU centroids.

#### 3.5.3 Phylogenetic analysis

Alignment of SSU rRNA genes was done with SSU-Align which is based on Infernal (Nawrocki *et al.*, 2009; Nawrocki and Eddy, 2013). Columns in the alignment that were not included in the SSU-Align covariance models or were aligned with poor confidence (less than 95% of characters in a position had posterior probability alignment scores of at least 95%) were masked for phylogenetic reconstruction. Additionally, the alignment was trimmed to coordinates such that all sequences in the alignment began and ended at the same positions. FastTree (Price *et al.*, 2010) was used to build the tree.

#### 3.5.4 Identifying OTUs that incorporated ^15^N into their DNA

DNA-SIP is a culture-independent approach towards defining identity-function connections in microbial communities (Radajewski and Murrell, 2001; Neufeld *et al.*, 2007; Buckley, 2011). Microbes are identified on the basis of isotope assimilation into DNA. As the buoyant density of a macromolecule is dependent on many factors in addition to stable isotope incorporation (e.g. G+C-content in nucleic acids (Youngblut and Buckley, 2014)), labeled nucleic acids from one microbial population may have the same buoyant density as unlabeled nucleic acids from another. Therefore, it is imperative to compare results of isotopic labelling to results obtained with unlabeled controls where everything mimics the experimental conditions except that unlabeled substrates are used. By contrasting heavy gradient fractions from isotopically labeled samples relative to corresponding fractions from controls, the identities of microbes with labeled nucleic acids can be determined

We used an RNA-Seq differential expression statistical framework (Love *et al.*, 2014) to find OTUs enriched in heavy fractions of labeled gradients relative to corresponding density fractions in control gradients (for review of RNA-Seq differential expression statistics applied to microbiome OTU count data see McMurdie and Holmes 2014). We use the term differential abundance (coined by McMurdie and Holmes 2014) to denote OTUs that have different proportion means across sample classes (in this case the only sample class is labeled:control). CsCl gradient fractions were categorized as “heavy” or “light”. The heavy category denotes fractions with density values above 1.725 g mL^−1^. Since we are only interested in enriched OTUs (labeled versus control), we used a one-sided Wald-test to test the statistical significance of regression coefficients (the null hypothesis is that the labeled:control fold enrichment for an OTU is less than a selected threshold). We independently filtered out sparse OTUs prior to P-value correction for multiple comparisons. The sparsity threshold was set to the value which maximized the number of p-values under a false discovery rate (FDR) the specific sparsity threshold was 0.3 meaning that an OTU not found in at least 30% of heavy fractions (control and labeled gradients) in a given day were not considered further and not included in P-value adjustment for multiple comparisons. P-values were corrected with the Benjamini-Hochberg method (Benjamini and Hochberg, 1995)and a FDR of 0.10 was applied (this rate is the typical FDR threshold adopted during RNASeq analysis). We selected a log_2_ fold change null threshold of 0.25 (or a labeled:control fold enrichment of 1.19). DESeq2 was used to calculate the moderated log_2_ fold change of labeled:control proportion means and corresponding standard errors for the Wald-test (above). Fold change moderation allows for reliable ranking such that high variance and likely statistically insignificant fold changes are appropriately shrunk and subsequently ranked lower than they would be as unmoderated values. Those OTUs that exhibit a statistically significant increase in proportion in heavy fractions from ^15^N_2_-labeled samples relative to corresponding controls have increased significantly in buoyant density in response to ^15^N_2_ treatment; a response that is expected for N_2_-fixing organisms.

We also assessed the consistency of enrichment between time points by including the interaction of day and label:control in a DESeq2 generalized linear model. The interpretation of the interaction coefficient is the change in OTU enrichment per unit time. P-values for the interaction coefficient were adjusted for all OTUs that passed the sparsity threshold in the label versus control comparison (above) and we used the default null model that the coefficient equaled zero. Additionally, we assessed fold change between labeled and control gradient heavy fractions after pooling day 2 and day 4 data when treating the different time points as replicates. The same null model as the label versus control comparison (above) was used in this replicate analysis (log_2_ fold change in abundance between label and control is less than or equal to 0.25). We included all OTUs that passed sparsity based independent filtering at either day (above) for p-value adjustment in the replicate analysis.

#### 3.5.5 Community and Sequence Analysis

BLAST searches were done with the “blastn” program from BLAST+ toolkit (Camacho *et al.*, 2009) version 2.2.29+. Default parameters were always employed and the BioPython (Cock *et al.*, 2009) BLAST+ wrapper was used to invoke the blastn program. Pandas (McKinney, 2012) and dplyr (Wickham and Francois, 2014) were used to parse and manipulate BLAST output tables.

Principal coordinate ordinations depict the relationship between samples at each time point (day 2 and 4). Bray-Curtis distances were used as the sample distance metric for ordination. The Phyloseq (McMurdie and Holmes, 2014) wrapper for Vegan (Oksanen *et al.*, 2013) (both R packages) was used to compute sample values along principal coordinate axes. GGplot2 (Wickham, 2009) was used to display sample points along the first and second principal axes. Adonis tests (Anderson, 2001) were done with default number of permutations (1000).

Rarefaction curves were created using bioinformatics modules in the PyCogent Python package (Knight *et al.*, 2007). Parametric richness estimates were made with CatchAll using only the best model for total OTU estimates (Bunge, 2010).

All code to take raw sequencing data through the presented figures (including download and processing of literature environmental datasets) can be found at: https://github.com/chuckpr/NSIP_data_analysis

## 4 RESULTS

### 4.1 DNA BUOYANT DENSITY CHANGES IN RESPONSE TO ^15^N_2_

BSCs were wetted and incubated for 4 days in transparent chambers with headspace containing N_2_ either from air or from 100 percent atom enriched ^15^N_2_. The chambers were illuminated with 16 h on / 8 h off cycles at an intensity of 200 *μ*mol photons m^−2^ s^−1^, which is the equivalent of an overcast/rainy day. N_2_-fixation as measured by acetylene reduction increased from 4.8 *μ*moles m^−2^ d^−1^ on day 2 to 14.5 *μ*moles m^−2^ d^−1^ on day 4. Amplicon sequences from ^15^N_2_-labeled samples and their corresponding unlabeled controls diverged specifically in heavy gradient fractions (Figure 1 and Figure S1) as assessed by Bray-Curtis dissimilarity (Bray and Curtis, 1957), and this result was significant (Adonis test (Anderson, 2001); p-value: 0.001, r^2^: 0.18).

**Figure 1.**
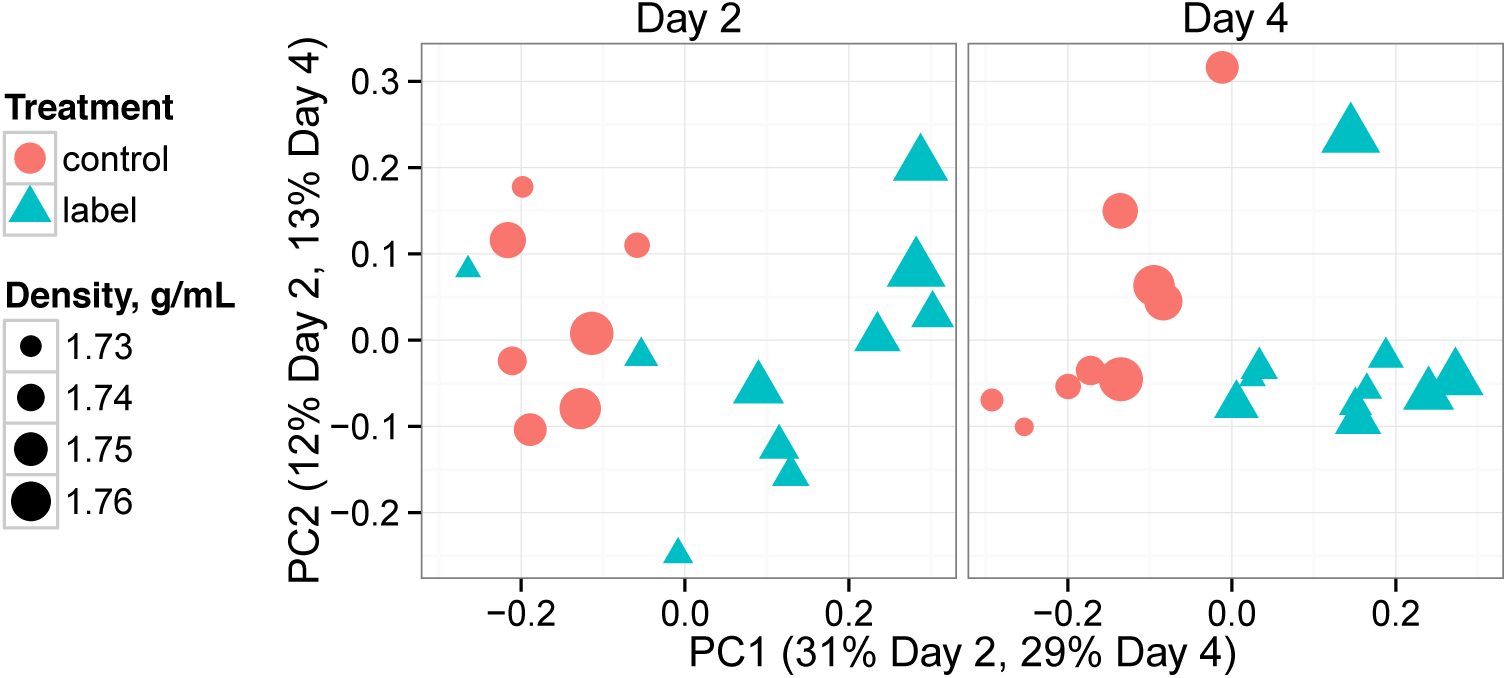
Ordination of heavy gradient fractions by Bray-Curtis distances on the basis of OTU content. Each point represents a gradient fraction OTU profile. Points closer together have more similar OTU content than those further apart.

### 4.2 OTUS RESPONSIVE TO ^15^N_2_ ARE PRIMARILY *PROTEOBACTERIA* AND *CLOSTRIDIACEAE*

OTUs that incorporated ^15^N into their DNA were detected by a differential change in their abundance within heavy gradient fractions of ^15^N_2_-labeled samples relative to corresponding controls. A total of 2,127 and 2,160 OTUs were detected in days 2 and 4, respectively, and these OTUs were interrogated for evidence of ^15^N_2_-labelling. Of these OTUs, only 499 and 563, respectively, passed a sparsity threshold applied to filter out OTUs with insufficient data for statistical analysis (see Love *et al.* (2014) for discussion of independent filtering). Of OTUs passing the sparsity criterion, 34 were enriched significantly in heavy fractions relative to control and this result is specifically expected for OTUs that have ^15^N-labeled DNA (*i.e.* ^15^N_2_ “responders”). Of these, 19 are annotated as *Firmicutes*, 12 as *Proteobacteria*, 2 as *Actinobacteria* and 1 as *Gemmatimonadetes* (Figure 2, Figure 3). If the responder OTUs are ranked by descending enrichment in heavy gradient fractions versus control, 8 the top 10 responders (*i.e.* those most enriched in the heavy fractions of labeled gradients) are either *Firmicutes* (3 OTUs) or *Proteobacteria* (5 OTUs) (Figure 4). Centroids (seed sequences) for strongly responding *Proteobacteria* OTUs all share high SSU rRNA gene sequence identity (>98.48%, Table 1) with isolates from genera known to possess diazotrophs including *Pseudomonas*, *Klebsiella*, *Shigella*, and *Ideonella*. None of the *Firmicutes* OTU centroids in the top 10 responders share greater than 97% SSU rRNA gene sequence identity with sequences in the Living Tree Project (LTP) database of 16S rRNA gene sequences from type strains (release 115) (see Table 1). OTUs that passed the sparsity threshold but were not classified as ^15^N-responsive were subsequently tested with the null hypothesis that the OTU fold enrichment in labeled gradient heavy fractions versus control was above the selected threshold. Rejecting the second null indicates that an OTU did *not* incorporate ^15^N into biomass. There were 86 and 89 “non-responders” at days 2 and 4, respectively. The ^15^N labelling of OTUs that did not pass sparsity or could not be classified as either a responder or non-responder cannot be determined conclusively.

**Figure 2.**
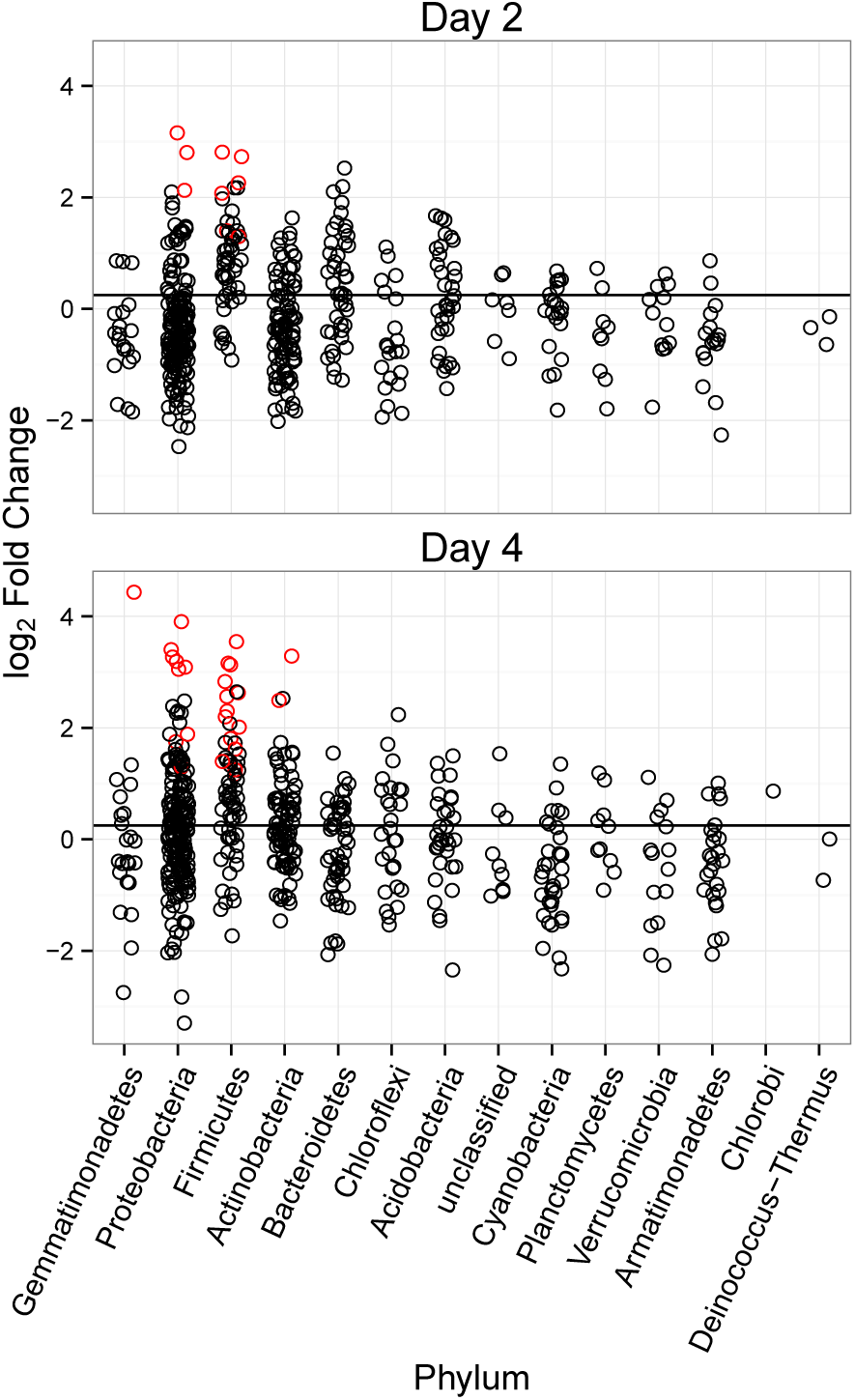
Moderated log_2_ fold change of OTUs proportions for labeled versus control gradients (heavy fractions only, densities >1.725 g/mL). All OTUs passing the sparsity treshold (see methods) at a specific incubation day are shown. Red color denotes a proportion fold change that has a corresponding adjusted p-value below a false discovery rate of 10% (ratio is significantly greater than 0.25, black line.)

**Figure 3.**
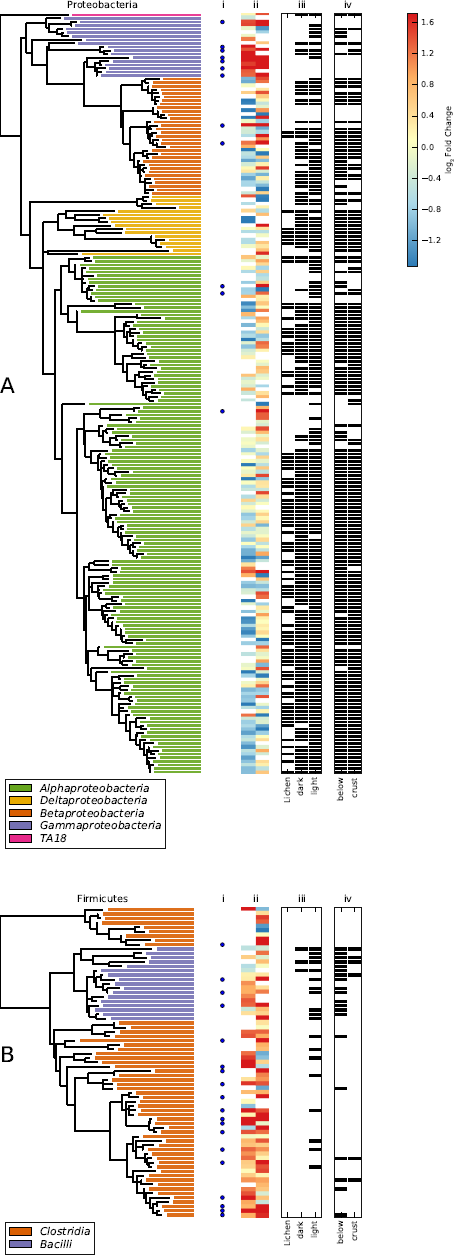
Phylogenetic trees of OTUs passing sparsity threshold for *Proteobacteria* **A** and *Firmicutes* **B**. ^15^N-responders are identified by dots present in column **i**. Log_2_ of OTU proportion fold change (labeled:control samples) for each OTU are presented as a heatmap in column **ii** with results from days 2 and 4 on the left and right sides of the column respectively. High fold change values indicate ^15^N incorporation. Presence/absence of OTUs (black indicates presence) in lichen, light, or dark environmental samples (Garcia-Pichel *et al.*, 2013) is shown in column **iii**. Presence/absence of OTUs (black indicates presence) in crust and below crust samples (Steven *et al.*, 2013) is shown in column **iv**.

**Figure 4.**
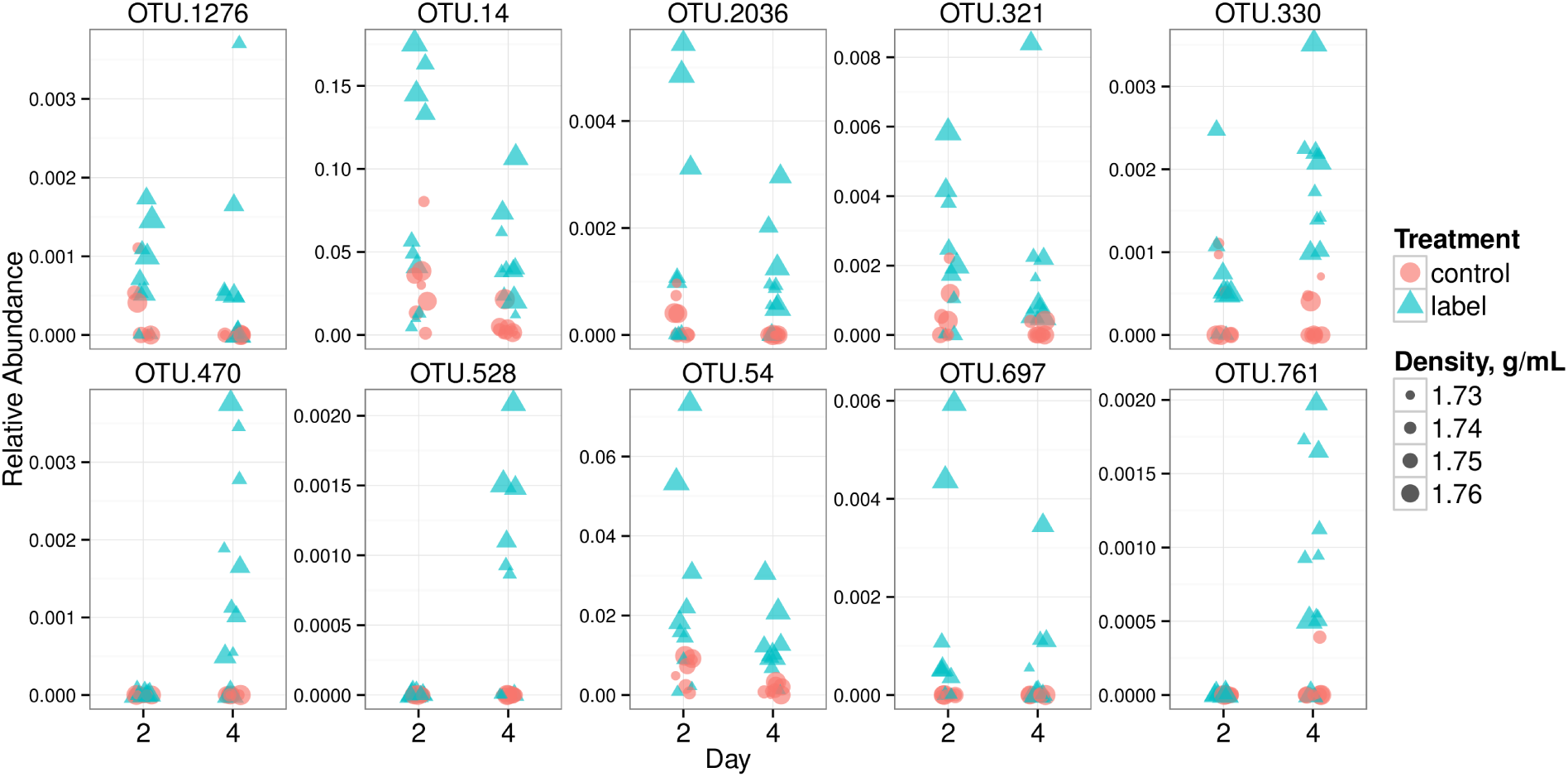
Relative abundance values in heavy fractions (density greater or equal to 1.725 g/mL) for the top 10 ^15^N “responders” (putative diazotrophs, see results for selection criteria of top 10) at each incubation day. Each point is a relative abundance value for the indicated OTU in a CsCl gradient fraction SSU rRNA gene collection. See Table 1 for BLAST results against the LTP database (release 115). Point area is proportional to CsCl gradient fraction density, and color signifies control (red) or labeled (blue) treatment.

**Table 1.** ^15^N responders BLAST search against Living Tree Project. Genera of all top BLAST hits are shown. Top 10 indicates responder was among top 10 most enriched OTUs in labeled gradient heavy fractions relative to corresponding control heavy fractions

| OTU ID | Genera | BLAST %ID | Top 10? | Phylum |
| --- | --- | --- | --- | --- |
| OTU.108 | *Caloramator* | 96.94 | no | *Firmicutes* |
| OTU.1276 | *Agromyces* | 99.49 | yes | *Actinobacteria* |
| OTU.137 | *Azospirillum* | 99.48 | no | *Proteobacteria* |
| OTU.14 | *Klebsiella, Kluyvera, Erwinia, Enterobacter, Pantoea, Buttiauxella* | 99.49 | yes | *Proteobacteria* |
| OTU.140 | *Bacillus* | 100.0 | no | *Firmicutes* |
| OTU.1673 | *Clostridium* | 95.9 | no | *Firmicutes* |
| OTU.176 | *Delftia* | 100.0 | no | *Proteobacteria* |
| OTU.2036 | *Pseudomonas* | 99.49 | yes | *Proteobacteria* |
| OTU.227 | *Cellulosilyticum* | 93.4 | no | *Firmicutes* |
| OTU.243 | *Bacillus* | 98.98 | no | *Firmicutes* |
| OTU.259 | *Parasporobacterium* | 98.47 | no | *Firmicutes* |
| OTU.263 | *Azospirillum* | 98.48 | no | *Proteobacteria* |
| OTU.278 | *Symbiobacterium* | 90.62 | no | *Firmicutes* |
| OTU.2794 | *Enterobacter* | 100.0 | no | *Proteobacteria* |
| OTU.282 | *Nocardia, Rhodococcus* | 100.0 | no | *Actinobacteria* |
| OTU.3 | *Bacillus* | 100.0 | no | *Firmicutes* |
| OTU.321 | *Pseudomonas* | 100.0 | yes | *Proteobacteria* |
| OTU.327 | *Clostridium* | 94.92 | no | *Firmicutes* |
| OTU.330 | *Clostridium* | 96.94 | yes | *Firmicutes* |
| OTU.342 | *Acinetobacter* | 100.0 | no | *Proteobacteria* |
| OTU.3712 | *Clostridium, Eubacterium* | 96.43 | no | *Firmicutes* |
| OTU.4037 | *Fonticella* | 93.85 | no | *Firmicutes* |
| OTU.4167 | *Fonticella* | 93.43 | no | *Firmicutes* |
| OTU.419 | *Caloramator* | 93.88 | no | *Firmicutes* |
| OTU.470 | *Gemmatimonas* | 85.86 | yes | *Gemmatimonadetes* |
| OTU.528 | *Clostridium* | 95.38 | yes | *Firmicutes* |
| OTU.54 | *Shigella, Escherichia* | 100.0 | yes | *Proteobacteria* |
| OTU.57 | *Fonticella, Caloramator* | 93.88 | no | *Firmicutes* |
| OTU.586 | *Ottowia, Diaphorobacter, Ideonella, Vitreoscilla, Comamonas* | 98.48 | no | *Proteobacteria* |
| OTU.61 | *Clostridium* | 95.92 | no | *Firmicutes* |
| OTU.643 | *Clostridium* | 97.45 | no | *Firmicutes* |
| OTU.647 | *Magnetospirillum* | 99.48 | no | *Proteobacteria* |

Table 1 – continued from previous page
| OTU ID | Genera | BLAST %ID | Top 10? | Phylum |
| --- | --- | --- | --- | --- |
| OTU.697 | *Pseudomonas* | 98.47 | yes | *Proteobacteria* |
| OTU.761 | *Gracilibacter* | 93.91 | yes | *Firmicutes* |

Although we did not take replicate samples within a time point we can assess the consistency of each OTUs response across the two time points. OTU fold enrichment at day 2 and 4 was consistent (Figure S2). There was a significant correlation between OTU fold enrichment at day 2 versus day 4 (P-value 4.35e^−8^). When the enrichment at day 2 is compared to day 4 via an interaction term (day *\** label/control, see methods), we found only two OTUs had significantly different enrichment between time points (”OTU.227” and “OTU.4037”, Table 1). In addition, when day 2 and day 4 samples are treated as replicates (see methods) only five of the OTUs we identified as responders OTUs were not significantly enriched in labeled gradient heavy fractions versus control (”OTU.140”, “OTU.4037”, “OTU.227”, “OTU.137”, and “OTU.263”, Table 1). The labeling of these OTUs should be interpreted with caution. None of the top 10 strongest responding OTUs showed inconsistent enrichment across time points based on the above analyses. Further, confidence in enrichment (i.e. lowest enrichment P-values between Day 2 and Day 4) appears to be correlated with consistency in response across both days (Figure S2).

### 4.3 ^15^N-RESPONSIVE OTUS ARE FOUND IN LOW ABUNDANCE IN AVAILABLE ENVIRONMENTAL BSC SSU RRNA GENE SURVEYS

In total 13 of the 34 ^15^N-responsive OTUs have been observed previously in SSU rRNA gene surveys of BSC communities (Figure 3, Figure S3). Eleven of the 19 ^15^N-responsive *Firmicutes* OTUs are members of the *Clostridiaceae*. Three ^15^N-responsive *Clostridiaceae* have been observed in previous BSC SSU rRNA gene surveys. Two ^15^N-responsive *Clostridiaceae* were found in “light” (*i.e.* early successional) crust during SSU rRNA gene sequence analysis of BSC (Garcia-Pichel *et al.*, 2013), and one ^15^N-responsive *Clostridiaceae* OTU was found among the “below crust” BSC SSU rRNA gene sequences described by Steven *et al.* (2013) (Figure 3). Five ^15^N-responsive proteobacterial OTUs (Table 1) were detected previously in BSC samples (Garcia-Pichel *et al.*, 2013; Steven *et al.*, 2013) The ^15^N-responsive *Gemmatimonadetes* OTU was observed in four Steven *et al.* (2013) samples and one ^15^N-responsive *Actinobacteria* OTU was found in three Steven *et al.* (2013) samples. *Gemmatimonadetes* and *Actinobacteria* ^15^N-responsive OTUs were not observed in samples collected by Garcia-Pichel *et al.* (2013)

### 4.4 COMPARISON OF SSU RRNA GENE SEQUENCES FROM DIFFERENT BSC SAMPLES

We compared the SSU rRNA gene sequences determined in this DNA-SIP experiment with two previous surveys of SSU rRNA gene amplicons from BSC communities (Garcia-Pichel *et al.*, 2013; Steven *et al.*, 2013). There were 3,079 OTUs (209,354 total sequences after quality control) in the DNA-SIP data, 3,203 OTUs (129,033 total sequences after quality control) in the Garcia-Pichel *et al.* (2013) study, and 2,481 OTUs (129,358 total sequences after quality control) in the Steven *et al.* (2013) study with a total of 4,340 OTUs across all three datasets. Of the total 4,340 OTU centroids established for this study, 445 have matches in the Living Tree Project (LTP) (a collection of SSU rRNA gene sequences for all sequenced type strains (Yarza *et al.*, 2008)) at or above a threshold of 97% sequence identity (LTP version 115). That is, 445 of 4,340 OTUs are closely related to known isolates. The DNA-SIP data shares 56% OTUs with the Steven *et al.* (2013) data and 46% of OTUs with the Garcia-Pichel *et al.* (2013) data, while these latter two studies share 46% of their OTUs. This result suggests that low frequency OTUs likely remain undersampled in all datasets.

Sequencing of DNA subjected to CsCl fractionation is expected to sample a different subset of diversity than that sampled by sequencing of unfractionated bulk DNA. For example, SIP enhances detection of OTUs that incorporate ^15^N into their DNA, and these OTUs will be overrepresented in the overall DNA-SIP sequence pool relative to their relative abundance in unfractionated bulk community samples. In addition, the DNA-SIP sequencing effort was directed at a relatively small number of “light” crust samples (n = 4), while previous sequencing efforts (Garcia-Pichel *et al.*, 2013; Steven *et al.*, 2013) were spread across hundreds of samples from both “light” and “dark” crusts. Hence, it is likely that the current study will be more likely to detect rare OTUs present in early successional “light” crust communities, particularly those that incorporate ^15^N into DNA. In all three BSC studies, most sequences were annotated as either cyanobacteria or *Proteobacteria*, though only in the DNA-SIP data did the sequences of *Proteobacteria* outnumber those of cyanobacteria. *Proteobacteria* represented 29.8% of sequence annotations in DNA-SIP data as opposed to 17.8% and 19.2% for the Garcia-Pichel *et al.* (2013) and Steven *et al.* (2013) data, respectively. In addition, sequences annotated as *Firmicutes* were more abundant in the DNA-SIP data (19%) than in the data from Steven *et al.* (2013) and Garcia-Pichel *et al.* (2013) (0.21% and 0.23%, respectively) (Figure S4). Finally, and congruently with sampling design sequences annotated to “Subsection IV” of cyanobacteria, which encompasses the heterocystous cyanobacteria in the Silva taxonomic nomenclature (Pruesse *et al.*, 2007), comprised only 0.29% of cyanobacteria sequences in the DNA-SIP data while representing 15% and 23% of cyanobacteria sequences from the Steven *et al.* (2013) and Garcia-Pichel *et al.* (2013) data, respectively.

## 5 DISCUSSION

BSC N-fixation has long been attributed to heterocystous cyanobacteria and the preponderance of cyanobacterial *nifH* genes observed in molecular surveys of BSCs have generally supported this hypothesis (Yeager *et al.*, 2004, 2007, 2012). However, in this study ^15^N_2_-DNA-SIP reveals that non-cyanobacterial microorganisms fix N_2_ in early successional BSC samples. *Proteobacteria* and *Clostridiaceae* were most abundant among ^15^N_2_-responsive OTUs as revealed by a robust statistical framework for quantifying and evaluating differential OTU abundance in microbiome studies (Love *et al.*, 2014; McMurdie and Holmes, 2014). Many of these OTUs (about 40%) have been observed previously in BSC communities. Rarefaction curves of data from Steven *et al.* (2013) and Garcia-Pichel *et al.* (2013) are still sharply increasing especially for sub-crust samples (Figure S5) suggesting the communities remain undersampled. Parametric richness estimates of BSC diversity indicate that the Steven *et al.* (2013) and Garcia-Pichel *et al.* (2013) sequencing efforts recovered on average 40.5% (s.d. 9.99%) and 45.5% (s.d. 11.6%) of predicted SSU rRNA gene OTUs from crust samples (inset Figure S5), respectively. Therefore, it would have been surprising if all of the ^15^N-responsive OTUs had been observed in prior environmental surveys of BSCs. Nitrogenase *nifH* gene sequences related to both *Proteobacteria* and *Clostridiaceae* have been previously observed in BSC samples, though typically at relative abundance that is much lower than *nifH* gene sequences from heterocystous cyanobacteria.

We propose three mechanisms that could bias *nifH* clone libraries against heterotrophic diazotrophs. First, extreme polyploidy in cyanobacteria (up to 58x ploidy in stationary phase,(Griese *et al.*, 2011)) can be expected to inflate the representation of cyanobacteria *nifH* gene sequences in community DNA relative to the frequency of ^15^N_2_-fixing heterocysts. Although, cyanobacteria often have relatively large cells so ploidy per cell is probably greater than ploidy per unit volume. Second, heterocysts make up a small fraction of total cells along a trichome, though all cells in the trichome possess the *nifH* gene. As a result of polyploidy and heterocyst frequency in a cyanobacterial filament, the ratio of cyanobacterial *nifH* gene copies to heterotrophic *nifH* gene copies may be inflated as much as 103 times relative to the corresponding ratio of ^15^N_2_-fixing cells (i.e. the ratio of heterocyst number to the cell number of heterotrophic diazotrophs). Third, *nifH* PCR primers, which are highly degenerate, could be biased against heterotrophic diazotrophs. For example, the *nifH* PCR primers used in the second round of a widely used nested PCR protocol (Yeager *et al.*, 2004, 2007, 2012) have fairly low coverage for *Proteobacteria* and *Clostridiales* (Gaby and Buckley, 2012). Primer “nifH11” is biased against “Cluster III” *nifH* gene sequences which includes those of the *Clostridiales* (50% *in silico* coverage of reference *nifH* sequences). In addition, primer “nifH22” has low coverage of reference sequences from *Proteobacteria*, cyanobacteria and “Cluster III” *nifH* gene sequences (16%, 23% and 21% *in silico* coverage, respectively) (Gaby and Buckley, 2012). Hence, it is reasonable to assume that heterotrophic diazotrophs may have been underestimated in previous analyses of early successional BSC communities. Our DNA-SIP results, which do not require PCR of functional genes, suggest that BSC N-fixation in early successional BSC may include a large non-cyanobacterial component. This is consistent with small-scale, spatially resolved functional measurements of nitrogen fixation in BSCs (Johnson *et al.*, 2005) that show a subsurface maximum that does not coincide spatially with maxima in chlorophyll *a* (a proxy for phototrophic biomass) in early-successional crusts, and a surface maximum of N_2_-fixation in mature crust that coincides with the maximum in chlorophyll *a*.

We did not observe incorporation of ^15^N_2_ into the DNA of heterocystous cyanobacteria in the early successional BSC samples used in this study. It is possible that ^15^N_2_-fixation by heterocystous cyanobacteria could go undetected in DNA-SIP. One possible explanation for this result is that the early successional BSC samples used in this study possessed too few heterocystous cyanobacteria to statistically evaluate their ^15^N-incorporation. Indeed, heterocystous cyanobacteria represented only 0.29% of sequences from the DNA-SIP data (see results) as opposed to 15% and 23% of total sequences in the Steven *et al.* (2013) and Garcia-Pichel *et al.* (2013) data, respectively. OTUs that correspond to heterocystous cyanobacteria (as defined by Yeager *et al.* (2007)), all fall below the sparsity threshold used in our analysis (see methods). Given the sparsity of heterocystous cyanobacteria sequences in the light crust DNA-SIP data, it is not possible to conclusively determine whether heterocystous cyanobacteria incorporated ^15^N during the incubation. Our results show that heterotrophic diazotrophs can contribute to ^15^N_2_-fixation in early successional BSC but they do not exclude the potential for fixation by heterocystous cyanobacteria. Indeed, heterocystous cyanobacteria if present, active, and limited for nitrogen would be expected to form heterocysts and fix ^15^N_2_. It is likely that scarcity limits their contribution to ^15^N_2_-fixation in early successional crusts. Heterocystous cyanobacteria form sessile colonies and they require stabilization of the crust environment before they can successfully colonize soil; and this stabilization is performed by other pioneering members of the crust community (Castenholz and Garcia-Pichel, 2002). ^15^N_2_-DNA-SIP would also fail to identify ^15^N_2_-fixing bacteria if ^15^N_2_-fixation were uncoupled from DNA replication over the time frame of the experiment (i.e. 4 days), that is ^15^N_2_-DNA-SIP will not detect bacteria that fix ^15^N_2_ but do not incorporate the ^15^N-label into DNA. Therefore, the contribution of heterocystous cyanobacteria (or any other microbe) to ^15^N_2_ would be underestimated if their cell division is uncoupled from ^15^N_2_-fixation at time frames of up to 4 days. We should also note that ^15^N can be incorporated into biomass from trophic interactions although in this case the ^15^N labeling would likely be weaker than that for a N_2_-fixer as a results of label dilution.

The OTUs with significant evidence of ^15^N-incorporation during the incubation were predominantly *Proteobacteria* and *Firmicutes*. The *Proteobacteria* OTUs with the strongest signal of ^15^N-incorporation all shared high sequence identity (>98.5%) with SSU rRNA gene sequences from genera known to contain diazotrophs (Table 1). In contrast the *Firmicutes* that displayed signal for ^15^N-incorporation (predominantly *Clostridiaceae*) were not closely related to any known cultivars (Table 1). Hence, we have little knowledge of the ecology of these organisms. Assessing the physiological characteristics of these diazotrophic *Clostridiaceae* may be useful for predicting how environmental change will affect the development and stability of BSC. Prior intense cultivation efforts from these crusts in separate studies did not yield any members of the *Clostridiaceae* (Gundlapally and Garcia-Pichel, 2006). Although under sampled in environmental data sets, ^15^N-responsive OTUs were indeed more abundant in sub-crust or in early successional BSC samples relative to crust surface or mature crust samples (Figure 3 and Figure S3). While members of *Clostridiaceae* have been found in low abundance in molecular surveys of BSC, most surveys are carried out on desicated crust samples, where thick-walled spores would predominate relative to vegetative cells, thus increasing the likelihood for their underrepresentation in DNA surveys. It should also be noted that crusts were incubated in an atmosphere of He and N_2_ rather than O_2_ and N_2_. While cyanobacteria in the presence of light rapidly produce oxygen super saturation in BSC relative to air (Garcia-Pichel and Belnap, 1996), and whereas heterotrophic N_2_-fixation by many microorganisms is inhibited in the presence of atmospheric levels of O_2_, it remains possible that the conditions present in microcosm are not representative of field conditions and may have favored N_2_-fixation by crust organisms that are less active in situ. Further experiments will need to be performed to verify that these heterotrophic diazotrophs are contributing to the N budgets of early successional crusts in the field.

Our results generate more refined hypotheses pertaining to the contribution of diazotrophs during the development of BSC communities. Specifically, ^15^N_2_-fixation in BSC may not be tied solely to the climax of heterocystous cyanobacteria in mature crusts. Rather, ^15^N_2_-fixation may occur throughout crust development with the transition between early successional and mature crusts marked by a transition between heterotrophic and phototrophic ^15^N_2_-fixation in the crust community. Therefore, sub-biocrust soil may contribute significantly to the arid ecosystem N budget and may be of considerable importance in the early phases of BSC establishment. We propose that interactions between fast-growing heterotrophic diazotrophs such as members of the *Clostridiaceae* and filamentous (non-heterocystous) cyanobacteria are important in the early establishment of BSC communities. During progressive desication, cyanobacteria, such as *M. vaginatus*, accumulate compatible solutes such as trehalose and sucrose (Rajeev *et al.*, 2013). Upon wetting, microorganisms rapidly excrete compatible solutes to prevent cell lysis due to osmotic shock (Poolman and Glaasker, 1998). Among them are dihexoses (such as sucrose and trehalose), which are observed in natural crusts upon wetting and then are rapidly depleted in the soil solution (Northen, 2014). Many *Clostridiaceae* have a saccharolytic metabolism with the potential for rapid growth rates on substrates such as trehalose and/or sucrose (Wiegel *et al.*, 2006). Wetting of crust may allow for rapid germination and growth of these organisms as the time required for germination of clostridial spores can be less than 30 minutes (Stringer *et al.*, 2005). Indeed, intense blooms of clostridia have been detected in crusts within tens of hours of wetting (Karaoz *et al.*, 2014). N_2_-fixing clostridia are common in soils (Wiegel *et al.*, 2006) and it is notable that *C. pasteurianum*, isolated from soil, was the first N_2_ fixing bacterium ever described (Winogradsky, 1895). *C. pasteurianum*, though an anaerobe, grows readily in the presence of oxygen when co-cultured with aerobic organisms that reduce oxygen tension (Chester, 1903). We propose that during a typical precipitation event, water saturation and heterotrophic activity rapidly render the interior of the crusts anoxic (Garcia-Pichel and Belnap, 1996) presenting optimal conditions for growth of anaerobic, dihexose-fermenting, N_2_ fixing clostridia. Clostridial organic nitrogen would then become available to other members of the community, including the primary producers, when carbon limitation induces sporulation and mother cell lysis. Mother cell lysis, the last step in sporulation, releases rich sources of P and N into the environment in the form of nucleotides and peptides (Hoch *et al.*, 2002).

### 5.1 CONCLUSION

The abundance of ^15^N-responsive OTUs from *Clostrideaceae* and *Proteobacteria* found in this study, the *nifH* gene sequences of *Clostrideaceae* and *Proteobacteria* observed previously in BSC (Steppe *et al.*, 1996), and the evidence for subsurface N_2_-fixation in early successional BSC (Johnson *et al.*, 2005), taken together, suggest that heterotrophic diazotrophs may be important contributors to N_2_-fixation in the subsurface of early successional BSC. Heterocystous cyanobacteria are also key contributors to the BSC N-budget, however and it is clear that heterocystous cyanobacteria increase in abundance with BSC age (Yeager *et al.*, 2004). It is less clear if the transition to mature crust is marked mainly by a change in the abundance and activity of heterocystous cyanobacteria, or rather represents a succession within the diazotroph community from early crusts where ^15^N_2_-fixation is dominated by *Clostridiaceae* and *Proteobacteria* to mature crusts where it is dominated by heterocystous cyanobacteria. Predicting the ecological response of BSC to climate change, altered precipitation regimes, and physical disturbance requires an understanding of crust establishment, stability, and succession. Diazotrophs are critical contributors to all of these phenomena and their activities make critical contributions to the N-budget of arid ecosystems worldwide.

## 6 ACKNOWLEDGEMENTS

We would like to thank T Whitman, CHD Williamson, AN Campbell and EK Hall for helpful comments in the preparation of this manuscript. This material is based upon work supported by the Department of Energy Office of Science, Office of Biological & Environmental Research Genomic Science Program under Award Numbers DE-SC0004486 and DE-SC0010558. This project was also supported by Agriculture and Food Research Initiative Competitive Grant no. 2007-35107-18299 from the USDA National Institute of Food and Agriculture. This report was prepared as an account of work sponsored by an agency of the United States Government. Neither the United States Government nor any agency thereof, nor any of their employees, makes any warranty, express or implied, or assumes any legal liability or responsibility for the accuracy, completeness, or usefulness of any information, apparatus, product, or process disclosed, or represents that its use would not infringe privately owned rights. Reference herein to any specific commercial product, process, or service by trade name, trademark, manufacturer, or otherwise does not necessarily constitute or imply its endorsement, recommendation, or favoring by the United States Government or any agency thereof. The views and opinions of authors expressed herein do not necessarily state or reflect those of the United States Government or any agency thereof.

## 7 CONFLICT OF INTEREST

The authors declare no conflict of interest.

## 10 SUPPLEMENTAL FIGURES

**Figure S1.**
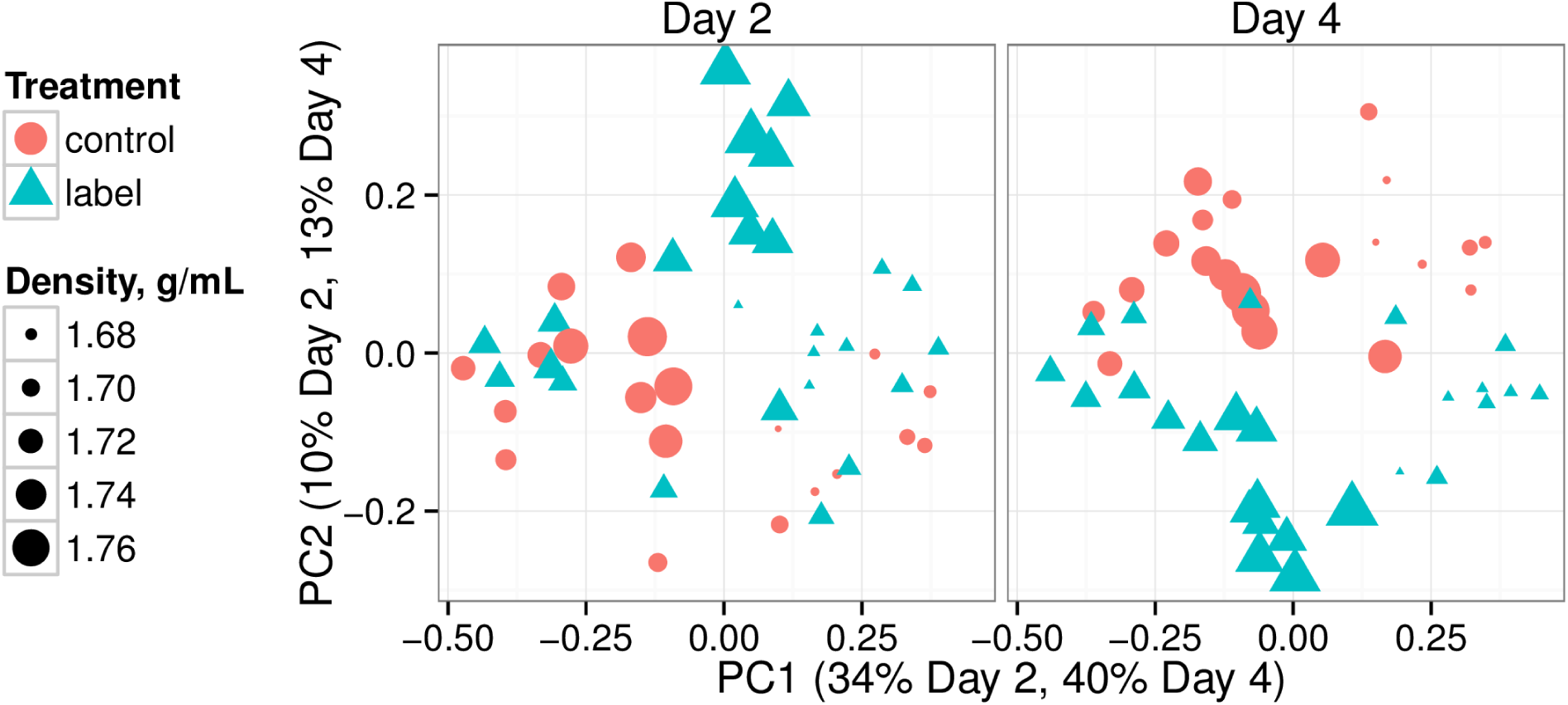
Ordination of Bray-Curtis sample pairwise distances for each incubation time. Point area is proportional to the density of the CsCl gradient fraction for each sequence library, and color/shape reflects control (red triangles) or labeled (blue circles) treatment. Each point represents the OTU profile for a single gradient fraction. Points closer together are more similar in OTU content than those further apart.

**Figure S2.**
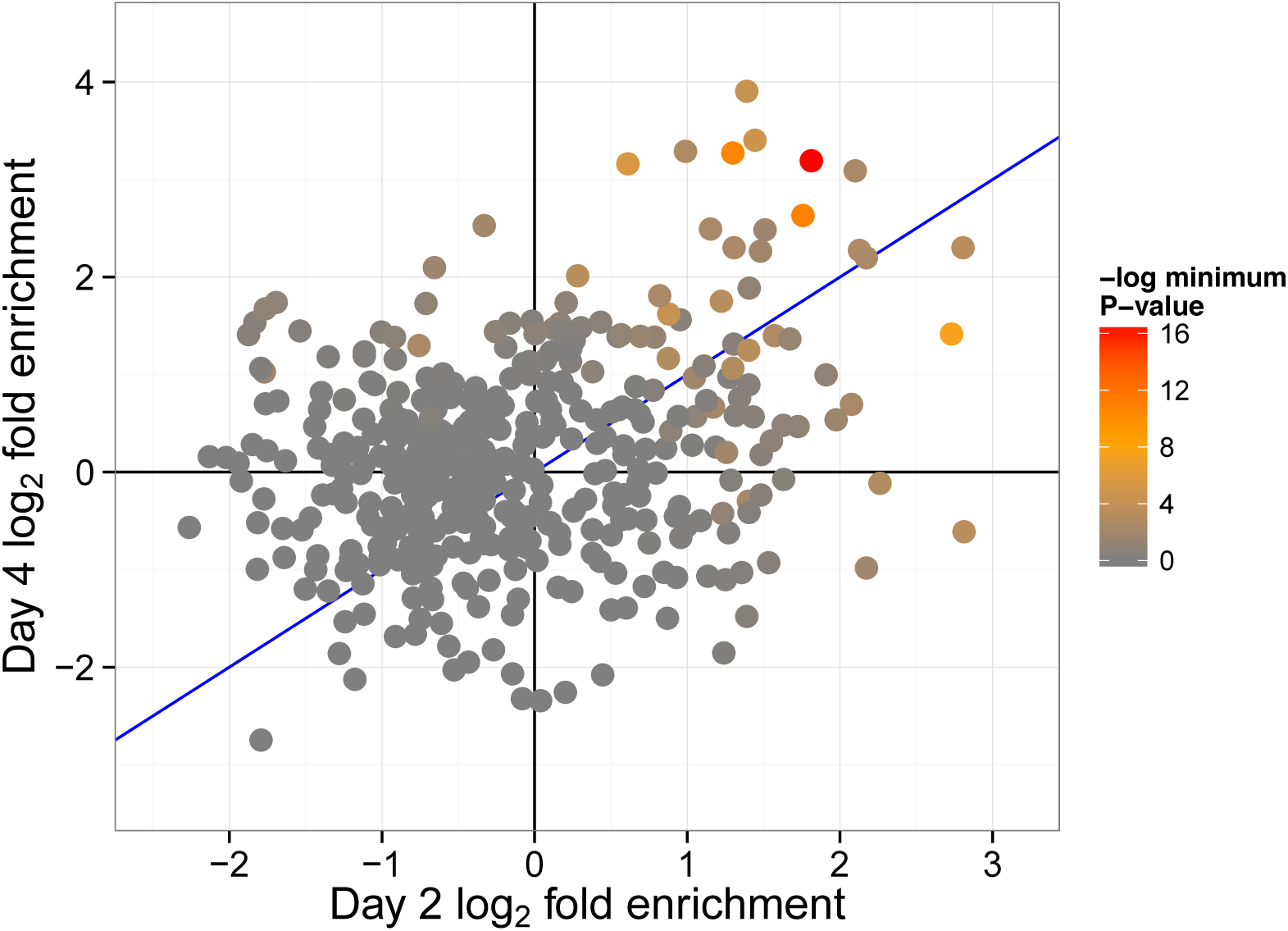
Scatter plot of fold enrichment values (label versus control heavy fractions) for OTUs passing sparsity criteria in day 2 and 4. P-value is the minimum value from day 2 or 4. Blue line has slope of one.

**Figure S3.**
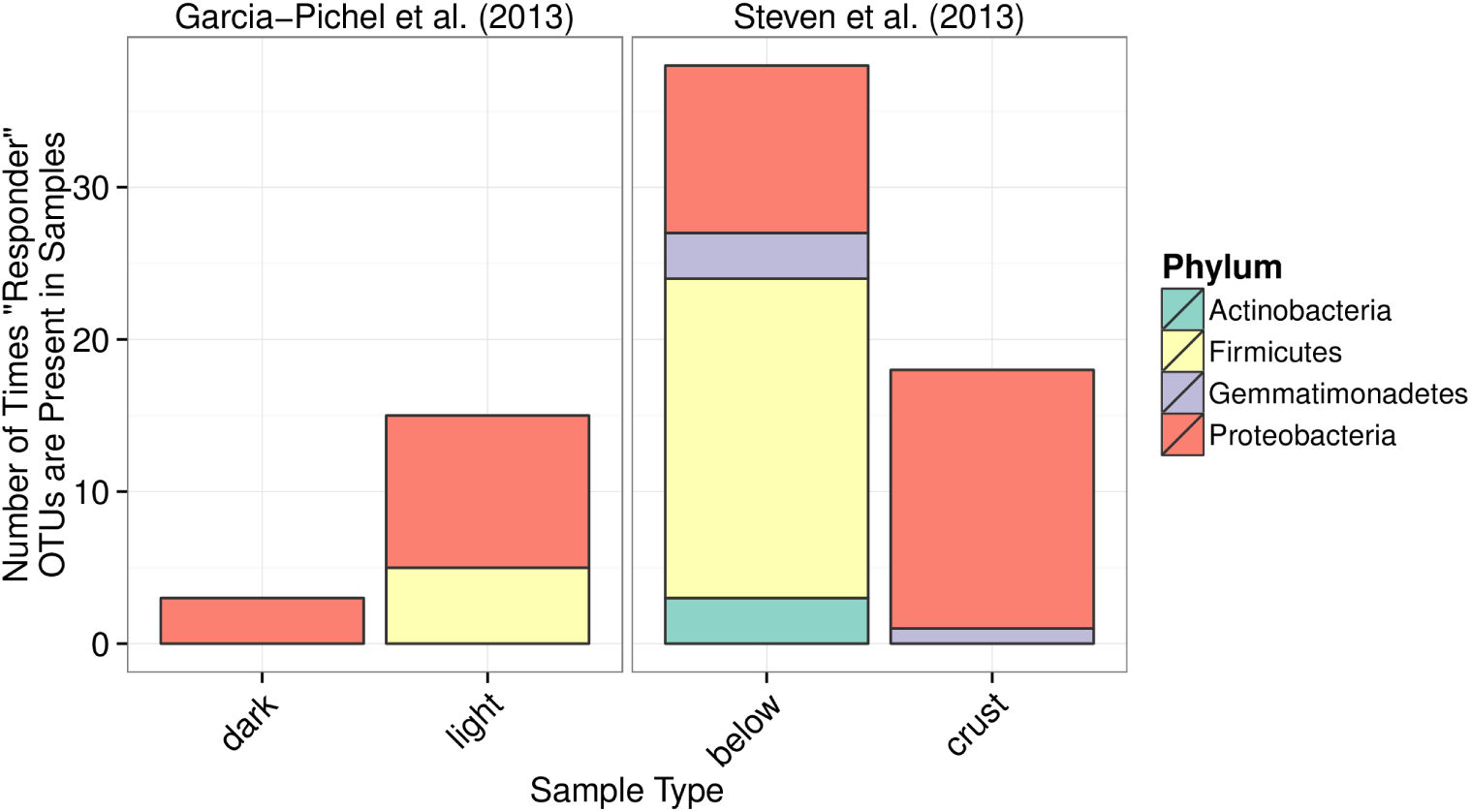
Counts of “responder” OTU occurrences in samples from Steven *et al.* (2013) and Garcia-Pichel *et al.* (2013) Steven *et al.* (2013) collected BSC samples (25 samples total) and samples from soil beneath BSC (17 samples total, “below” column in figure). Garcia-Pichel *et al.* (2013) collected samples from “dark” (9 samples total) and “light” (12 samples total) crusts in addition to “Lichen” (2 samples total) dominated crusts.

**Figure S4.**
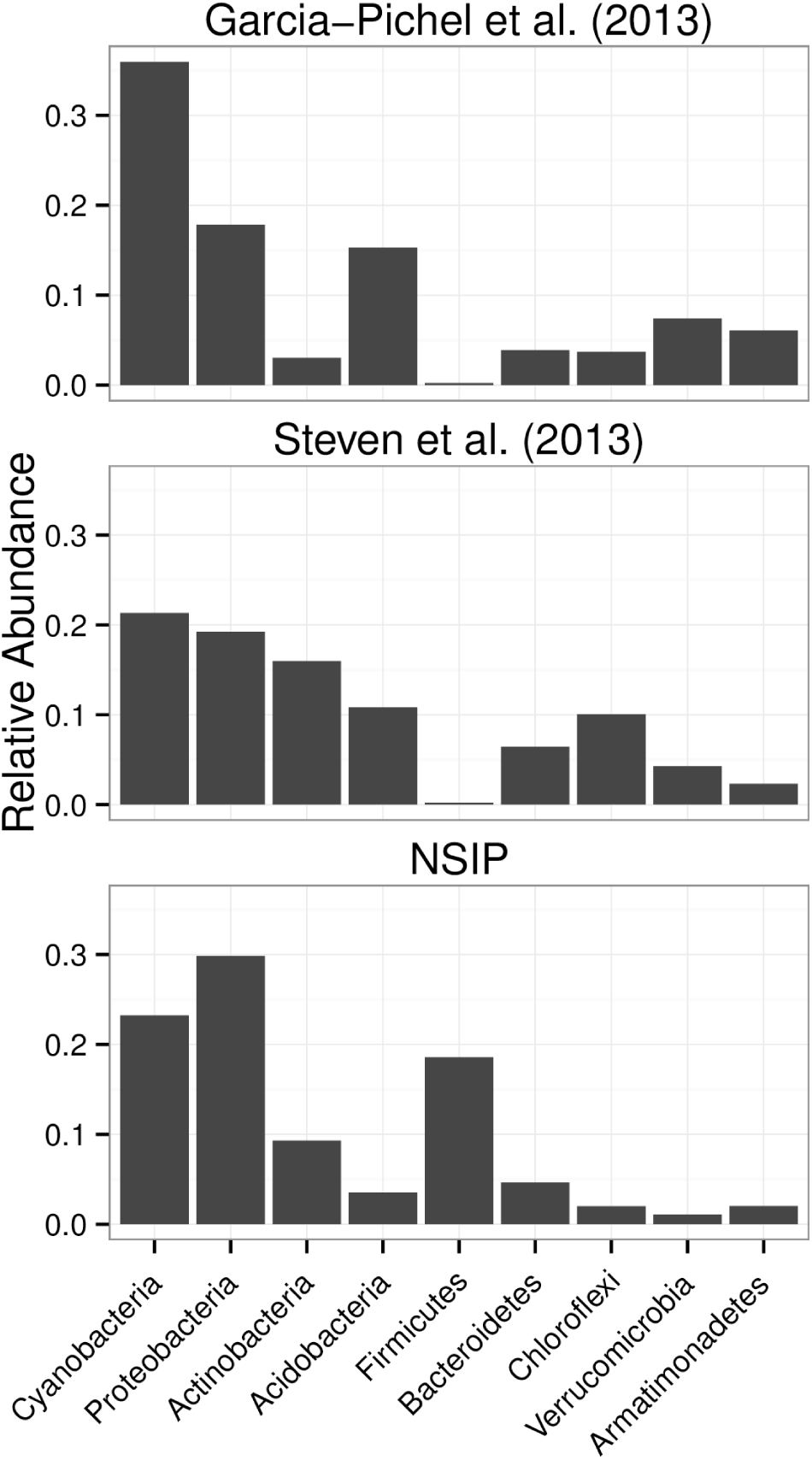
Distribution of sequences into top 9 phyla (phyla ranked by sum of all sequence annotations).

**Figure S5.**
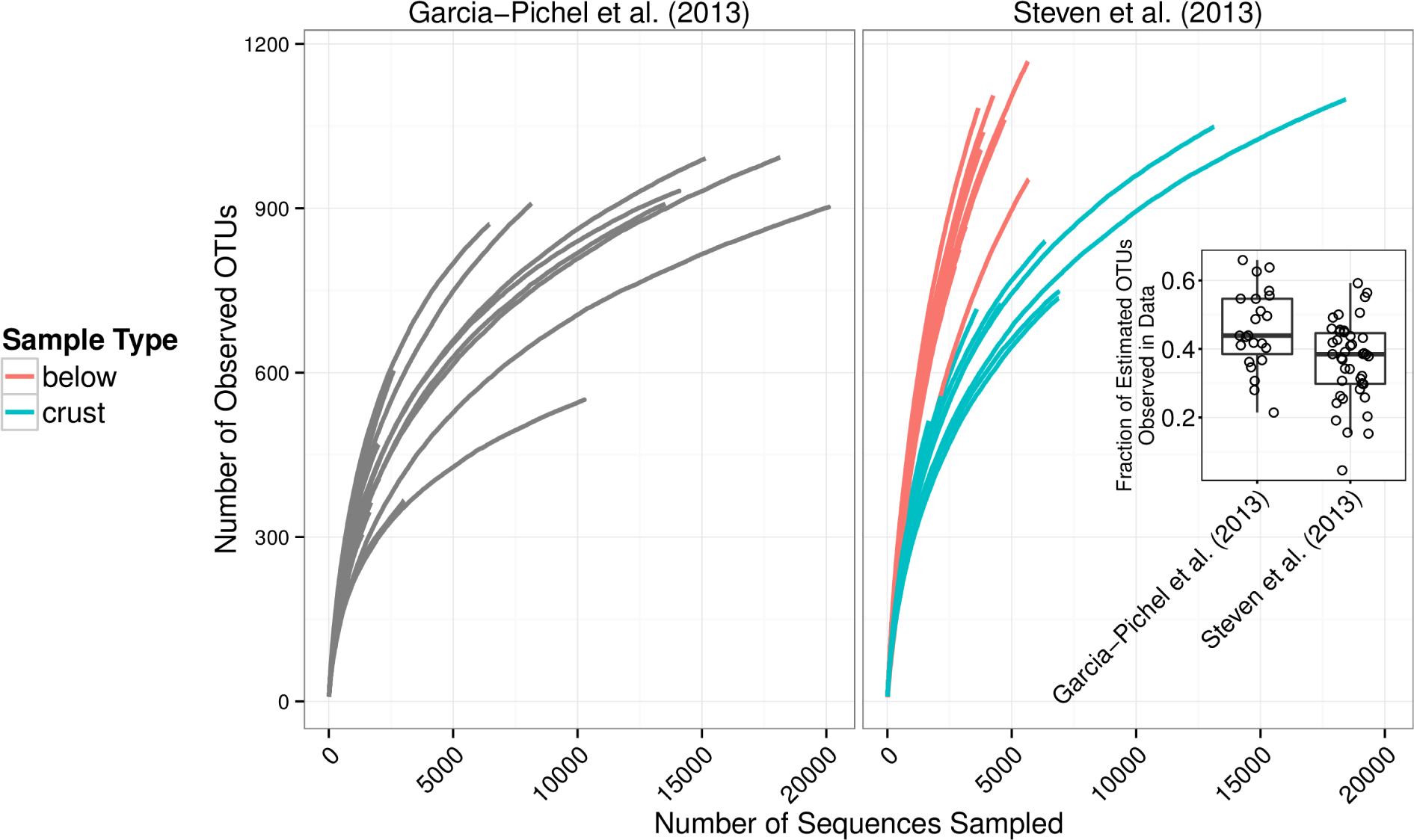
Rarefaction curves for all samples presented by Garcia-Pichel *et al.* (2013) and Steven *et al.* (2013) Inset is boxplot of estimated sampling effort for all samples in Garcia-Pichel *et al.* (2013) and Steven *et al.* (2013) (number of observed OTUs divided by number of CatchAll (Bunge, 2010) estimated total OTUs)

## REFERENCES

Anderson M. (2001). A new method for non-parametric multivariate analysis of variance. Austral Ecology 26: 32–46.

Belnap J. (2001). Factors influencing nitrogen fixation and nitrogen release in biological soil crusts. In: Belnap J, Lange O (eds.) Biological soil crusts: structure, function, and management. Vol. 150. Ecological Studies. Springer: Berlin Heidelberg, pp. 241–261.

Belnap J. (2002). Nitrogen fixation in biological soil crusts from southeast Utah USA Biol Fert Soils 35: 128–135.

Belnap J. (2003). Factors influencing nitrogen fixation and nitrogen release in biological soil crusts. In: Belnap J, Lange O (eds.) Biological soil crusts: structure, function, and management. Vol. 150. Ecological Studies. Springer: Berlin Heidelberg, pp. 241–261.

Belnap J, Prasse R, Harper K. (2003). Influence of biological soil crusts on soil environments and vascular plants. In: Belnap J, Lange O (eds.) Biological soil crusts: structure, function, and management. Vol. 150. Ecological Studies. Springer: Berlin Heidelberg, pp. 281–300.

Benjamini Y, Hochberg Y. (1995). Controlling the false discovery rate: a practical and powerful approach to multiple testing. J R Stat Soc Series B Stat Methodol 57: 289–300.

Beraldi-Campesi H, Hartnett H, Anbar A, Gordon G, Garcia-Pichel F. (2009). Effect of biological soil crusts on soil elemental concentrations: implications for biogeochemistry and as traceable biosignatures of ancient life on land. Geobiology 7: 348–359.

Bray J, Curtis J. (1957). An ordination of the upland forest communities of southern Wisconsin. Ecol Monograph 27:325.

Buckley D. (2011). Stable isotope probing techniques using ^15^N In: Murrell J, Whiteley A (eds.) Stable isotope probing and related technologies. American Society of Microbiology Press: Washington. DC pp. 129–147.

Buckley D, Huangyutitham V, Hsu S, Nelson T. (2007). Stable isotope probing with ^15^N_2_ reveals novel noncultivated diazotrophs in soil. Appl Environ Microbiol 73: 3196–3204.

Bunge J. (2010). Estimating the number of species with Catchall. In: Altman R, Dunker, L H, Murray T, Klein T (eds.) Biocomputing 2011. World Scientific: Hackensack, NJ pp. 121–130.

Camacho C, Coulouris G, Avagyan V, Ma N, Papadopoulos J, Bealer K et al.. (2009). BLAST+: Architecture and applications. BMC Bioinformatics 10: 421.

Caporaso J, Kuczynski J, Stombaugh J, Bittinger K, Bushman F, Costello E et al.. (2010). QIIME allows analysis of high-throughput community sequencing data. Nat Methods 7: 335– 336.

Castenholz RW, Garcia-Pichel F. (2002). Cyanobacterial Responses to UV-radiation. In: Whitton B, Potts M (eds.) The ecology of cyanobacteria. Springer: Netherlands, pp. 591– 611.

Chester F. (1903). Oligonitrophilic bacteria of the soil. Science. 370–371.

Cock P, Antao T, Chang J, Chapman B, Cox C, Dalke A et al.. (2009). Biopython: Freely available Python tools for computational molecular biology and bioinformatics. Bioinformatics 25: 1422–1423.

DeSantis TJ, Hugenholtz P, Keller K, Brodie E, Larsen N, Piceno Y et al.. (2006). NAST: a multiple sequence alignment server for comparative analysis of 16S rRNA genes. Nucleic Acids Res 34: W394–W399.

Edgar R. (2010). Search and clustering orders of magnitude faster than BLAST. Bioinformatics 26: 2460–2461.

Edgar R. (2013). UPARSE: highly accurate OTU sequences from microbial amplicon reads. Nat Methods 10: 996–998.

Evans R, Belnap J. (1999). Long-term consequences of disturbance on nitrogen dynamics in an arid ecosystem. Ecology 80: 150–160.

Evans R, Lange O. (2001). Biological soil crusts and ecosystem nitrogen and carbon dynamics. In: Belnap J, Lange O (eds.) Biological soil crusts: structure, function, and management. Vol. 150. Ecological Studies. Springer: Berlin Heidelberg, pp. 263–279.

Gaby J, Buckley D. (2012). A comprehensive evaluation of PCR primers to amplify the *nifH* Gene of nitrogenase. PLoS ONE 7: e42149.

Garcia-Pichel F, Loza V, Marusenko Y, Mateo P, Potrafka R. (2013). Temperature drives the continental-scale distribution of key microbes in topsoil communities. Science 340: 1574– 1577.

Garcia-Pichel F, Belnap J, Neuer S, Schanz F. (2003). Estimates of global cyanobacterial biomass and its distribution. Algol Stud 109: 213–227.

Garcia-Pichel F, Belnap J. (1996). Microenvironments and microscale productivity of cyanobacterial desert crusts. J Phycol 32: 774–782.

Garcia-Pichel F, Wojciechowski MF. (2009). The evolution of a capacity to build supra-cellular ropes enabled filamentous cyanobacteria to colonize highly erodible substrates. PLoS ONE 4: e7801.

Griese M, Lange C, Soppa J. (2011). Ploidy in cyanobacteria. FEMS Microbiol Lett 323: 124– 131.

Gundlapally SR, Garcia-Pichel F. (2006). The community and phylogenetic diversity of biological soil crusts in the colorado plateau studied by molecular fingerprinting and intensive cultivation. Microb Ecol 52: 345–357.

Hoch J, Sonenshein A, Losick R. (2002). Bacillus subtilis: From cells to genes and from genes to cells. In: Sonenshein A, Hoch J, Losick R (eds.) Bacillus subtilis and its closest relatives. American Society of Microbiology: Washington, DC

Johnson SL, Neuer S, Garcia-Pichel F. (2007). Export of nitrogenous compounds due to incomplete cycling within biological soil crusts of arid lands. Environ Microbiol 9: 680– 689.

Johnson S, Budinoff C, Belnap J, Garcia-Pichel F. (2005). Relevance of ammonium oxidation within biological soil crust communities. Environ Microbiol 7: 1–12.

Karaoz U, Estelle C, Nunes Da Rocha U, Northen T, Garcia-Pichel F, Brodie E. (2014). in preparation.

Karnieli A, Kokaly R, West N, Clark R. (2003). Remote sensing of biological soil crusts. In: Belnap J, Lange O (eds.) Biological soil crusts: structure, function, and management. Vol. 150. Ecological Studies. Springer: Berlin Heidelberg, pp. 431–455.

Knight R, Maxwell P, Birmingham A, Carnes J, Caporaso J, Easton B et al.. (2007). PyCogent: A toolkit for making sense from sequence. Genome Biol 8: R171.

Love MI, Huber W, Anders S. (2014). Moderated estimation of fold change and dispersion for RNA-Seq data with DESeq2. bioRxiv.

McKinney W. (2012). pandas: Python data analysis library. Online. URL: http://pandas.pydata.org/.

McMurdie P, Holmes S. (2014). Waste not, want not: why rarefying microbiome data is inadmissible. PLoS Comput Biology 10: e1003531.

Nawrocki E, Eddy S. (2013). Infernal 1.1: 100-fold faster RNA homology searches. Bioinformatics 29: 2933–2935.

Nawrocki E, Kolbe D, Eddy S. (2009). Infernal 1.0: inference of RNA alignments. Bioinformatics 25: 1335–1337.

Neufeld J, Vohra J, Dumont M, Lueders T, Manefield M, Friedrich M et al.. (2007). DNA stable-isotope probing. Nat Protoc 2: 860–866.

Northen T. (2014). personal communication.

Oksanen J, Blanchet F, Kindt R, Legendre P, Minchia nP, O’Hara R et al.. (2013). vegan: Community ecology package. R package version 2.0-10. URL: http://CRAN.R-project.org/package=vegan.

Poolman B, Glaasker E. (1998). Regulation of compatible solute accumulation in bacteria. Mol Microbiol 29: 397–407.

Price M, Dehal P, Arkin A. (2010). FastTree 2–approximately maximum-likelihood trees for large alignments. PLoS One 5: e9490.

Pruesse E, Quast C, Knittel K, Fuchs B, Ludwig W, Peplies J et al.. (2007). SILVA: a comprehensive online resource for quality checked and aligned ribosomal RNA sequence data compatible with ARB. Nucleic Acids Res 35: 7188–7196.

Radajewski S, Murrell J. (2001). Stable isotope probing for detection of methanotrophs after enrichment with 13CH4. In: Gene probes. Humana Press: New York, pp. 149–157.

Rajeev L, Rocha UN da, Klitgord N, Luning EG, Fortney J, Axen SD et al.. (2013). Dynamic cyanobacterial response to hydration and dehydration in a desert biological soil crust. ISME J 7: 2178–2191.

Schloss P, Westcott S, Ryabin T, Hall J, Hartmann M, Hollister E et al.. (2009). Introducing Mothur: open-source, platform-independent, community-supported software for describing and comparing microbial communities. Appl Environ Microbiol 75: 7537–7541.

Starkenburg SR, Reitenga KG, Freitas T, Johnson S, Chain PSG, Garcia-Pichel F et al.. (2011). Genome of the cyanobacterium *Microcoleus vaginatus* FGP-2 a photosynthetic ecosystem engineer of arid land soil biocrusts worldwide. J Bacteriol 193: 4569–4570.

Steppe T, Olson J, Paerl H, Litaker R, Belnap J. (1996). Consortial N_2_ fixation: a strategy for meeting nitrogen requirements of marine and terrestrial cyanobacterial mats. FEMS Microbiol Ecol 21: 149–156.

Steven B, Gallegos-Graves L, Belnap J, Kuske C. (2013). Dryland soil microbial communities display spatial biogeographic patterns associated with soil depth and soil parent material. FEMS Microbiol Ecol 86: 101–113.

Strauss SL, Day TA, Garcia-Pichel F. (2011). Nitrogen cycling in desert biological soil crusts across biogeographic regions in the southwestern united states. Biogeochem 108: 171–182.

Stringer SC, Webb MD, George SM, Pin C, Peck MW. (2005). Heterogeneity of times required for germination and outgrowth from single spores of nonproteolytic *Clostridium botulinum*. Appl Environ Microbiol 71: 4998–5003.

Walters W, Caporaso J, Lauber C, Berg-Lyons D, Fierer N, Knight R. (2011). PrimerProspector: de novo design and taxonomic analysis of barcoded polymerase chain reaction primers. Bioinformatics 27: 1159–1161.

Wickham H. (2009). ggplot2: elegant graphics for data analysis. Springer: New York.

Wickham H, Francois R. (2014). dplyr: dplyr: a grammar of data manipulation. R package. URL: http://CRAN.R-project.org/package=dplyr.

Wiegel J, Tanner R, Rainey F. (2006). An introduction to the family clostridiaceae. In: Rosenberg E, DeLong E, Lory S, Stackebrandt E, Thompson F (eds.) The prokaryotes. Springer: US pp. 654–678.

Winogradsky S. (1895). Recherches sur l’assimilation de l’azote libre de l’atmosphere par les microbes. Arch. d. Sci. Biol. 4: 297.

Yarza P, Richter M, Peplies J, Euzeby J, Amann R, Schleifer K et al.. (2008). The All-Species Living Tree project: A 16S rRNA-based phylogenetic tree of all sequenced type strains. Syst Appl Microbiol 31: 241–250.

Yeager C, Kornosky J, Housman D, Grote E, Belnap J, Kuske C. (2004). Diazotrophic community structure and function in two successional stages of biological soil crusts from the Colorado Plateau and Chihuahuan Desert. Appl Environ Microbiol 70: 973–983.

Yeager C, Kornosky J, Morgan R, Cain E, Garcia-Pichel F, Housman D et al.. (2007). Three distinct clades of cultured heterocystous cyanobacteria constitute the dominant N_2_-fixing members of biological soil crusts of the Colorado Plateau USA FEMS Microbiol Ecol 60: 85–97.

Yeager C, Kuske C, Carney T, Johnson S, Ticknor L, Belnap J. (2012). Response of biological soil crust diazotrophs to season altered summer precipitation, and year-round increased temperature in an arid grassland of the Colorado Plateau, USA Front Microbiol 3:

Youngblut ND, Buckley DH. (2014). Intra-genomic variation in G+C content and its implications for dna stable isotope probing. Environ Microbiol Reports 6: 767–775.

